# Counting Cases, Conserving Species: Addressing Highly Pathogenic Avian Influenza in Wildlife

**DOI:** 10.1101/2025.06.18.660293

**Authors:** Ulrich Knief, Sandra Bouwhuis, Anja Globig, Anne Günther, Wouter Courtens

**Affiliations:** University of Freiburg, Institute of Biology I (Zoology), Evolutionary Biology and Ecology, Hauptstr. 1, DE-79104 Freiburg, Germany; Institute of Avian Research, An der Vogelwarte 21, D-26386 Wilhelmshaven, Germany; Friedrich-Loeffler-Institut, Institute of International Animal Health/One Health, Federal Research Institute for Animal Health, Südufer 10, DE-17493 Greifswald-Insel Riems, Germany; Friedrich-Loeffler-Institut, Institute of Diagnostic Virology, Federal Research Institute for Animal Health, Südufer 10, DE-17493 Greifswald-Insel Riems, Germany; Research Institute for Nature and Forest, Havenlaan 88, bus 73, 1000 Brussels, Belgium

**Keywords:** HPAI, H5N1, wildlife, wild birds, ecology, mass mortality, disease outbreak

## Abstract

Highly pathogenic avian influenza (HPAI) has become a critical threat to wildlife, shifting from a seasonal epizootic to a persistent, year-round panzootic with global consequences. Here, we summarize the origin, evolutionary mechanisms, and expanding host range of the current H5N1 virus (clade 2.3.4.4b) and assess its impact on wildlife. Over the past five years, HPAI has caused the deaths of millions of wild birds, causing dramatic population declines in several seabird species. However, comprehensive quantitative mortality data remain scarce, as existing records are often anecdotal, focus on localized mass die-offs, and thus represent only a fraction of the true magnitude of mortality. This gap in data limits the ability to predict outbreak dynamics and mitigate long-term consequences. Using the Northwestern European Sandwich Tern (*Thalasseus sandvicensis*) population as a case study, we demonstrate the value of integrating mortality data with ecological, serological and genetic data before, during and after an outbreak. This approach uncovered age-specific vulnerability, selective mortality, and population immunological responses. In addition, insights gained with respect to the role of breeding density, carcass removal, and host adaptation in modulating outbreak dynamics are likely to be generalizable across seabird species. The absence of a centralized and standardized wildlife mortality monitoring framework, on the other hand, remains a major barrier to effective outbreak forecasting and conservation planning. We argue that integrating field-based mortality data, population monitoring, serological assays, and genetic analyses within a One Health framework is essential to enable early detection, targeted mitigation, and robust evaluation of outbreak impacts. Without a proactive and data-driven approach to conservation, HPAI will continue to threaten global wildlife populations, with cascading ecological, economic and public health consequences.

## 1. Origin and Properties of the Highly Pathogenic H5 Avian Influenza Virus Goose/Guandong 1996

The highly pathogenic avian influenza (HPAI) virus subtype H5N1 of clade 2.3.4.4b has made global headlines, causing the deaths of hundreds of millions of domestic poultry and wild animals (**Fig. 1**, **Supplement**; Weston 2024). Since 2021, its enzootic circulation in Europe (Pohlmann *et al*. 2022), expansion into new host species (Klaassen & Wille 2023), and transcontinental spread to the Americas (Caliendo *et al*. 2022) and Antarctica (Bennett-Laso *et al*. 2024) have left, and continue to leave, a trail of devastation around the world. HPAI, also commonly known as bird flu, is a highly contagious disease and often manifests with severe neurological symptoms, leading to high mortality in its avian hosts. Originally confined to poultry (“fowl plague”), HPAI has, in recent years, spilled over into a broad range of hosts, including wild birds, wild carnivorous and marine mammals, fur animals, pets, and even livestock.

**Figure 1.**
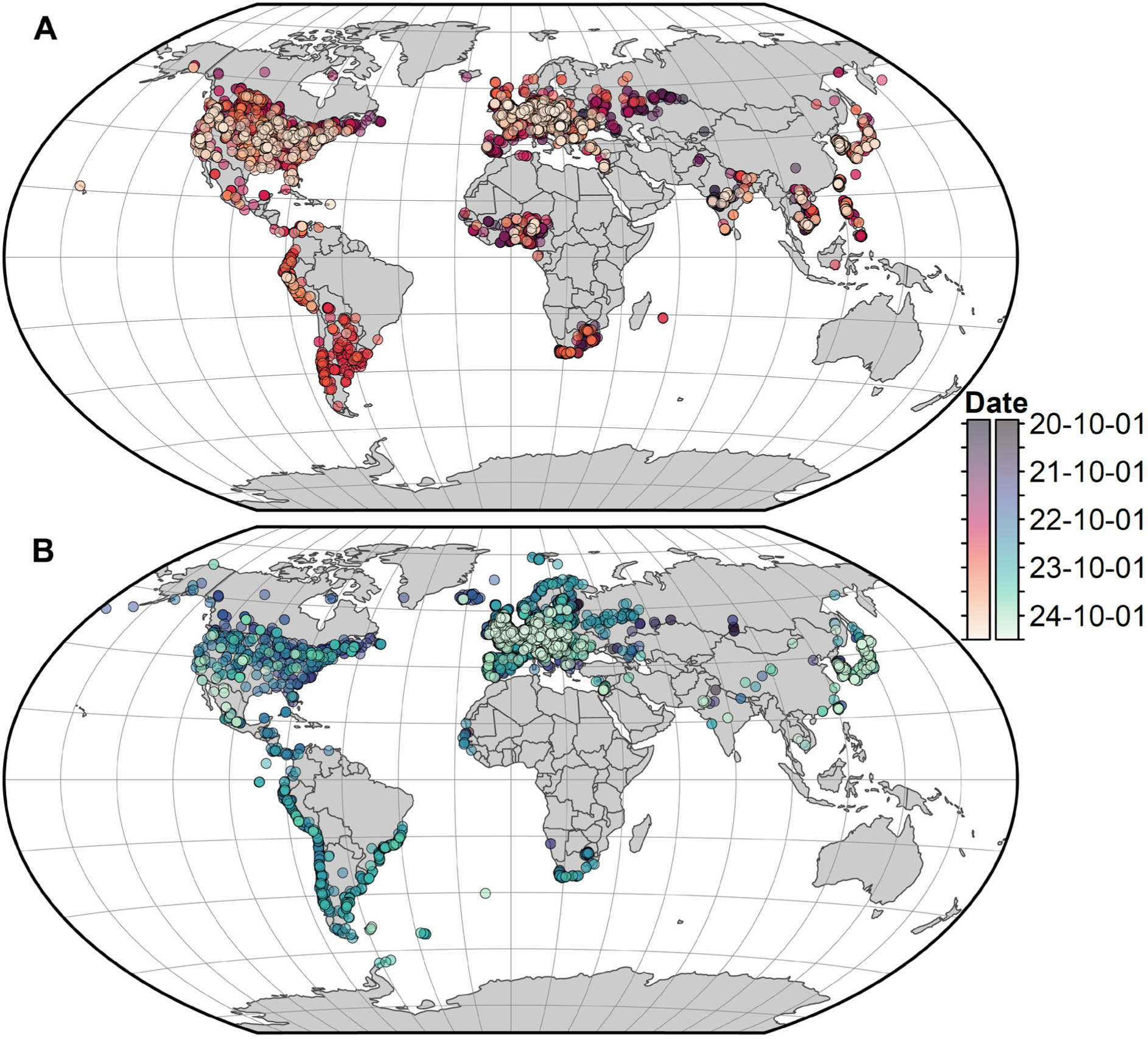
Spatiotemporal dynamics of the current HPAI H5N1 panzootic in (A) domestic (*N* = 9,078) and (B) wild (*N* = 9,388) animals. Each dot represents an avian influenza positive case from the EMPRES-i+ database (disease: “Influenza – Avian”) reported between 1 September 2020 and 1 April 2025. Dots may represent isolated cases or mass mortality events. The panzootic originated in Europe (index case October 2020 in the Netherlands; Lewis *et al*. 2021) and subsequently spread to Africa (December 2020; Abolnik *et al*. 2023) and East Asia (October 2021; Tian *et al*. 2023). Earlier occurrences in South and East Asia were of independent origin (Tare *et al*. 2024; Xie *et al*. 2023). North America was reached via two independent incursions: from Europe (November 2021) and East Asia (February 2022; Giacinti *et al*. 2024). The virus spread from North to South America for the first time in October 2022, likely via migratory birds, with initial outbreaks in Colombia (Leguia *et al*. 2023). From there, it moved clockwise around the continent, but also moved overland from west to east throughout 2022–2023 (Kuiken *et al*. 2025; Uhart *et al*. 2024). The virus reached the (sub)Antarctic islands: Falkland Islands in October 2023 (Banyard *et al*. 2024), South Georgia in September 2023 (Bennison *et al*. 2024), St. Helena and Marion Island in September 2024 (DFFE 2025), and Crozet and the Kerguelen Islands in October 2024 (not yet included in the EMPRES-i+ database; Clessin *et al*. 2025). Finally, it reached Antarctica in February 2024 (Bennett-Laso *et al*. 2024). Even remote oceanic archipelagos such as the Galápagos (September 2023; Stokstad 2023) and the Hawaiian Islands (November 2024; State of Hawai’i 2025) have been affected.

Until the 21^st^ century, HPAI infections were rare events, even in poultry (Alexander & Brown 2009). HPAI viruses emerge from low pathogenic avian influenza (LPAI) viruses of the H5 or H7 subtypes through mutations in the hemagglutinin (HA) gene (Graziosi *et al*. 2024). LPAI viruses typically cause asymptomatic infections and have long been circulating in wild and captive bird populations, where they show low rates of evolutionary change (Fourment & Holmes 2015). Mutations that convert LPAI into HPAI viruses are most likely to occur in environments with high bird densities, such as intensive poultry farms, where avian influenza viruses (AIVs) can easily replicate in susceptible hosts. Indeed, 95% of the known mutation events transforming LPAI into HPAI have occurred in captive poultry (Dhingra *et al*. 2018), after which the virus spilled over into populations of (migratory) wild birds and moved across the globe (Liu *et al*. 2005). As such, human agricultural practices are largely responsible for the current panzootic (Kuiken & Cromie 2022).

A defining characteristic of AIVs is their RNA-based genome, which is divided into eight segments. RNA mutates more readily than DNA (antigenic drift), and when a host is co-infected with two genetically distinct AIV subtypes, these segments can additionally exchange through a process called reassortment (antigenic shift), generating new virus variants with altered properties, driving the extraordinary variability and continuous evolution of AIVs (Dhingra *et al*. 2018). These changes may enable AIVs to evade host immune responses, through processes known as antigenic drift and antigenic shift (Luczo & Spackman 2024; Webster *et al*. 1992).

Although AIVs frequently reassort and exchange genetic material, it is the hemagglutinin gene that plays the central role in determining whether a virus exhibits high or low pathogenicity. Consequently, the evolutionary history of the HA gene is closely monitored, and its genetic divergence forms the basis for defining viral clades and subclades (Smith & Donis 2015).

The currently dominating HPAI viruses carry an H5 hemagglutinin gene and descends from a lineage first isolated from domestic geese in Guangdong, China, in 1996—later designated as Goose (Gs)/Guandong (Gd)/1/1996 (Alexander & Brown 2009). Initially confined to poultry in southeast China, it was detected in wild birds for the first time in 2003, and subsequently spread to waterfowl in multiple countries across Southeast Asia (Alexander 2007). Following its detection in wild migratory birds at Qinghai Lake in northwestern China in 2005 (Chen *et al*. 2005; Liu *et al*. 2005), the virus—later designated as clade 2.2 (Ducatez *et al*. 2017)—spread rapidly westward, reaching Europe, the Middle East, and Africa by 2006, with some evidence that wild migratory water birds contributed to this rapid spread (Global Consortium for H5N8 and Related Influenza Viruses 2016; Lee *et al*. 2016; Salzberg *et al*. 2007; Shi *et al*. 2023).

After this expansion, outbreaks in wild birds declined markedly, as clade 2.2 was largely eliminated in Asia and Europe within two years (King *et al*. 2021). In poultry in Egypt, however, it circulated for many years until being replaced by higher-order subclades, especially after 2016 with 2.3.4.4b (El-Shesheny *et al*. 2021; Shi *et al*. 2023). Continued circulation in domestic poultry, especially in Southeast Asia, facilitated further viral evolution (Sonnberg *et al*. 2013). Around 2005, clade 2.3.4 emerged in China (de Vries *et al*. 2015), and subsequently diversified into several subclades between 2008 and 2010 (Lee *et al*. 2016).

Among these, subclade 2.3.4.4 was first detected in 2010 (Global Consortium for H5N8 and Related Influenza Viruses 2016; Xie *et al*. 2023) and later identified in multiple countries across Asia, Europe and Africa (Antigua *et al*. 2019). Subclade 2.3.4.4 viruses are characterized by frequent reassortment, also described as “promiscuous”, thereby acquiring different neuraminidase (NA) gene segments (N1, N2, N5, N6, N8) from cocirculating LPAI viruses (de Vries *et al*. 2015; Global Consortium for H5N8 and Related Influenza Viruses 2016; Nguyen *et al*. 2025; Pohlmann *et al*. 2022). In late 2014, a 2.3.4.4 virus—later designated as subclade 2.3.4.4c (Graziosi *et al*. 2024)—was detected in North America for the first time, likely following intercontinental spread via migratory birds through the Bering Strait to Alaska (Global Consortium for H5N8 and Related Influenza Viruses 2016; Lee *et al*. 2015; Shi *et al*. 2023). However, its impact on wild birds remained limited, and it disappeared from North America by 2016 (Xie *et al*. 2023).

Subclade 2.3.4.4b was first detected in domestic ducks in eastern China in 2013 and in South Korea in 2014 (Graziosi *et al*. 2024 and references therein). A reassortant H5N8 virus of this subclade was isolated from wild birds during a mass mortality event at Qinghai Lake in 2016 (Li *et al*. 2017). This reassortant virus marked a turning point: it spread from China to Europe in 2016 and 2017, expanded to an unparalleled geographic range, affected a broader spectrum of species, and caused the largest documented outbreak in wild birds up to that point (King *et al*. 2021; Xie *et al*. 2023). The subsequent outbreak of 2020–2021 originated from a H5N8 virus that likely evolved in Egypt, and was even more severe (Xie *et al*. 2023). The current HPAI virus subtype H5N1 of clade 2.3.4.4b originated mid-2020 as a reassortant of this H5N8 virus and is likely of European origin. During this reassortment, the virus exchanged its N8 neuraminidase gene segment with an N1 segment that had been circulating in European wild birds since 2019, and it additionally replaced five internal gene segments (Pohlmann *et al*. 2022; Xie *et al*. 2023). This reassortant H5N1 virus was first detected in the Netherlands in October 2020 (called the “index case”) (Lewis *et al*. 2021). It is this virus that caused the ongoing outbreak, which has surpassed all previous events in geographic scale, ecological severity, and economic impact— spreading across continents and severely affecting wild bird populations and an unprecedented range of other wild animal species worldwide (Peacock *et al*. 2025).

## 2. An Overview of HPAI in Wildlife Since 2021 in Europe and Beyond

Before 2021, HPAI was a seasonal epizootic disease in Europe, primarily affecting domestic poultry and wild (land- and) waterfowl (*Galloanserae*), with occasional secondary transmission to raptors feeding on infected wild birds. Outbreaks were most pronounced in winter, such as among Barnacle Geese (*Branta leucopsis*) along the North Sea coast (King *et al*. 2021). Occasionally, shorebirds were affected as well, as seen in the Wadden Sea during the winter of 2020, when approximately 3,500 Red Knots (*Calidris canutus*) died (King *et al*. 2024).

### A Shift to Year-Round Infections and Global Spread

In 2021, the situation shifted dramatically. HPAI became enzootic in Europe (Pohlmann *et al*. 2022), meaning that while it was previously absent during summer, cases and outbreaks now occurred year-round in wild birds. They became particularly intense during the breeding season, especially in colonially-breeding seabirds, but also in secondary hosts (Günther *et al*. 2024; Nemeth *et al*. 2023). In 2022, for example, colonially-breeding Northern Gannets (*Morus bassanus*; Lane *et al*. 2024), Sandwich Terns (*Thalasseus sandvicensis*; Knief *et al*. 2024) and Common Terns (*Sterna hirundo*; Pohlmann *et al*. 2023) suffered substantial losses while breeding across the North and Baltic Seas. While the precise mutations or reassortments of the viral genome that enabled the initial expansion of host species remain unclear (or may not have occurred at all), a notable reassortment event in spring 2022 involved the circulating HPAI H5N1 and a gull-adapted LPAI, leading to the emergence of a new H5N1 genotype (EA-2022-BB) in northern France (Briand *et al*. 2025; EFSA *et al*. 2023b; Fusaro *et al*. 2024). This EA-2022-BB genotype appeared to be particularly adapted to gulls and terns, equally fatal to its hosts, highly contagious, and possibly transmitted through airborne routes (Nagy *et al*. 2025; Zhang *et al*. 2022). It spread among Black-headed Gulls (*Chroicocephalus ridibundus*) during the winter, and in the subsequent summer affected breeding colonies of Black-headed Gulls and Common Terns, from the coasts to the foothills of the Alps (Knief *et al*. in preparation).

Globally, HPAI viruses of clade 2.3.4.4b have now spread to every continent except Australia (Wille *et al*. 2024). HPAI reached the Americas in September 2021 (**Fig. 1**), and the most severe outbreaks among wild bird populations in North America occurred in eastern Canada during 2022, with approximately 40,391 recorded deaths, primarily among Northern Gannets and Common Murre (*Uria aalge*) (Avery-Gomm *et al*. 2024) (**Fig. 2**, **Supplement**). Subsequently, South America faced severe impacts in 2022 and 2023, when the virus affected immunologically naïve wild bird populations with no prior exposure to HPAI. An estimated 667,000 wild birds succumbed (Kuiken *et al*. 2025; Uhart *et al*. 2024) (**Fig. 2**, **Supplement**). Reports from Peru alone included 254,793 Guanay Cormorants (*Leucocarbo bougainvilliorum*) and Neotropical Cormorants (*Nannopterum brasilianus*), 235,643 Peruvian Boobies (*Sula variegata*), and 57,447 Peruvian Pelicans (*Pelecanus thagus*). This mortality represents up to 50% of the global populations of these species (Gamarra-Toledo *et al*. 2023; Kuiken *et al*. 2025).

**Figure 2.**
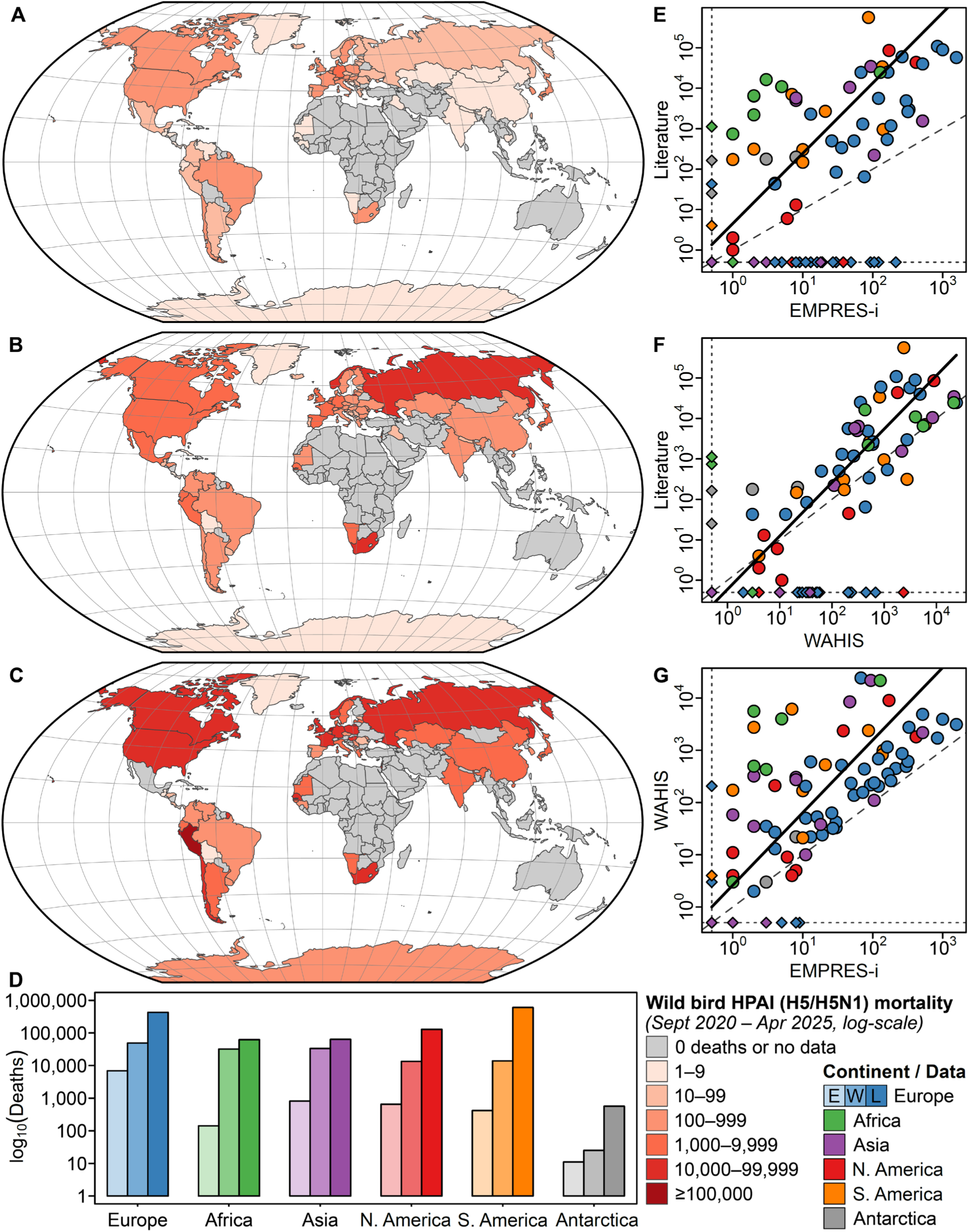
Comparison of wild bird mortality data (log-scale) from the EMPRES-i+ database, the World Animal Health Information System (WAHIS) database, and a literature review of reported mass mortality events since the emergence of HPAI subtype H5N1 clade 2.3.4.4b in autumn 2020 until April 2024. For this figure, the number of reported deaths per country was summed across species and age classes, as this information is often missing from databases and even from some published reports. (A–C) Number of reported deaths per country in (A) EMPRES-i+, (B) WAHIS, and (C) the literature review. (D) Total number of reported deaths per continent (left to right: E = EMPRES-i+, W = WAHIS, L = Literature). Overall, 8,928, 141,651, and 1,285,036 dead wild birds were reported through EMPRES-i+, WAHIS, and the literature, respectively. In addition, 460, 380, and 54,740 dead wild mammals were reported through EMPRES-i+, WAHIS, and the literature (not shown). (E–G) Pairwise comparisons between country-based datasets. Diamonds along horizontal and vertical dotted lines indicate missing records in one or the other dataset. These were excluded from the major axis regression. The dashed line represents the 1:1 line (perfect agreement), and the thick black line shows the major axis regression. Major axis regression slopes in (E) and (F) are significantly different from 1 (*P* = 5.4 × 10^-3^ and *P* = 7.8 × 10^-3^, respectively). Colours represent continents.

HPAI viruses reached the (Sub-)Antarctic region via South America in the summer of 2023, with confirmed outbreaks in the Falkland Islands and South Georgia (Banyard *et al*. 2024; Bennison *et al*. 2024). By February 2024, the virus had also been detected on the Antarctic mainland, marking the first known incursion of HPAI into the continent (Bennett-Laso *et al*. 2024). Later that year, it spread from South America to Prince Edward, Crozet, and Kerguelen Islands—remote archipelagos in the Southern Indian Ocean. Genetic analyses confirmed this eastward transatlantic transmission route, rather than spread via Africa (Clessin *et al*. 2025; Duvenage 2024). In autumn 2024, the virus also reached Hawai‘i in the Central Pacific, most likely carried by migratory birds from Alaska (State of Hawai’i 2025), and was detected on St. Helena in the Central Atlantic (EMPRES-i), further illustrating the global dispersal of the virus.

### Beyond Birds: HPAI Infections in Mammals

The global spread of HPAI has also led to increasing infections in wild mammals. While media attention has mostly focused on infected dairy cattle in North America (e.g. Capua & Fanelli 2024; Nguyen *et al*. 2025), carnivorous mammals have been most commonly affected in the wild. These animals likely contract the virus by consuming infected dead or weakened birds. Both carnivorous marine mammals, such as seals, porpoises, and dolphins, as well as terrestrial ones, including foxes, bears, and mustelids, have been affected in large numbers (ENETWILD Consortium *et al*. 2024; Plaza *et al*. 2024a; Puryear & Runstadler 2024). While outbreaks in terrestrial species mostly involved single, isolated cases, the outbreaks among marine mammals, particularly in South America, were especially severe. Between late 2022 and 2023, the virus spread along the coasts of Peru, Chile, Argentina, and Uruguay, eventually reaching Brazil. During this period, at least 31,894 South American Sea Lions (*Otaria flavescens*; Kuiken *et al*. 2025; Plaza *et al*. 2024b; Uhart *et al*. 2024) and 17,400 Southern Elephant Seals (*Mirounga leonina*) succumbed to the disease (Campagna *et al*. 2024; Kuiken *et al*. 2025). In Argentina’s Valdés Peninsula, the outbreak led to the death of more than half of the local Southern Elephant Seal population. Concerningly, during these mammalian outbreaks, the virus acquired mutations that enhanced its adaptation to mammals and seemingly enabled mammal-to-mammal transmission (Nguyen *et al*. 2025; Tomás *et al*. 2024; Uhart *et al*. 2024).

These quantitative wildlife data should not give the false impression that we have a comprehensive understanding of the dynamics and impact of HPAI outbreaks in wild animal populations (Couty *et al*. 2025; Klaassen & Wille 2023). In reality, they represent only a small fraction of what actually occurred, and a centralized system for collecting quantitative mortality data is urgently needed to estimate the true impact of HPAI in wild birds (see **Fig. 2**). In the following paragraphs, we illustrate what we can derive from such data and the questions that remain to be addressed. Using the HPAI outbreaks in European Sandwich Tern colonies as an example, and by drawing on additional relevant information, we try to generalize these findings to other species. We also highlight how these scientific endeavours can benefit species conservation, the economy, and public health. Finally, we propose concrete political actions that should be taken to improve our current understanding of HPAI in wildlife.

## 3. The Case of HPAI in Northwestern European Sandwich Terns

### Risk Factors for Sandwich Terns

Similar to many other seabirds, Sandwich Terns breed in dense colonies on the ground, with up to seven nests per square meter, spaced just far enough apart to prevent neighbours from reaching one another. These colonies can host several thousand breeding pairs, but there were only 67 colonies across Northwestern Europe in 2022, making Sandwich Terns an ideal model species to study HPAI dynamics both at a population and a meta-population level. The high breeding density, combined with the species’ tendency to defecate around nest sites, can result in elevated viral loads within colonies and facilitate both direct bird-to-bird and indirect transmission via faecal-oral routes or environmental contact. Regular inter- and intra-annual exchange of birds between colonies (Spaans *et al*. 2021, Van Bemmelen *et al*. in preparation) then creates a scenario in which, once the virus enters a colony and is sufficiently adapted to the host species, it can spread rapidly, potentially wiping out a significant proportion of the entire Northwestern European population.

### The 2022 Outbreak: Devastating Impact

The Northwestern European Sandwich Tern population suffered greatly from HPAI in 2022. Knief *et al*. (2024) collected data on breeding pair numbers, mortality rates, and whether carcasses were removed from colonies. Before the outbreak, the population comprised 59,632 breeding pairs, equivalent to 119,264 adult individuals, excluding late breeders and non-breeding birds such as prospectors. Most of the 65 colonies with HPAI status data were concentrated along the North Sea coasts of France, Belgium, the Netherlands, Germany, Denmark, and the United Kingdom. During the summer of 2022, over 19,326 adult birds were confirmed dead across 39 colonies in Northwestern Europe. Based on additional data, we estimated that total mortality exceeded 17% of the breeding population, amounting to more than 20,000 adult birds. Additionally, nearly all chicks in infected colonies perished. While occasional reproductive failure is less critical for this long-lived species, which is capable of surviving up to 32 years and adapted to low breeding success, the loss of adult birds poses a far greater threat to the population’s long-term viability (Sæther *et al*. 2016).

### Trends in 2023 and Beyond: Fewer Deaths and Evidence for an Immune Response

During the winter of 2022–2023, reports from West Africa, where Sandwich Terns overwinter, documented over 25,000 deaths among seabirds, including at least 2,394 Sandwich Terns (Barlow *et al*. 2023; Jatta *et al*. 2025; Ndao 2023).

Anticipating another devastating outbreak during the breeding season of 2023, the situation unfolded differently than expected. Breeding pair numbers fell by 25% to 44,748 pairs (Courtens *et al*. in preparation) (**Fig. 3**). However, while the virus remained present in at least 32 colonies, spread to colonies in the Mediterranean Sea (Valle *et al*. 2024) and caused another near-total loss of chicks in affected colonies, adult birds were largely spared. Only 592 adult Sandwich Terns were found dead, which represents a reduction in mortality of about 96% (**Fig. 3**). This unexpected shift raised questions. Although carcass removal had proven to be an effective mitigation measure (Knief *et al*. 2024) and was widely implemented in 2023, this alone could not explain the difference. The virus did not appear to change in virulence or contagiousness among gulls and terns (genotypes EA-2021-AB [terns] and EA-2020-C [Northern Gannets] were replaced with EA-2022-BB, which seemed to be even more gull-adapted) (Briand *et al*. 2025; Fusaro *et al*. 2024; Pohlmann *et al*. 2023), as it continued to heavily impact Black-headed Gulls and Common Terns, decimating colonies of these two species across Europe. Moreover, Sandwich Tern chicks still died in large numbers from EA-2022-BB in 2023 (all 41 genotyped individuals; EFSA *et al*. 2023a; Fusaro *et al*. 2024, Fusaro personal communication). So why did adult mortality decline after the initial outbreak?

**Figure 3.**
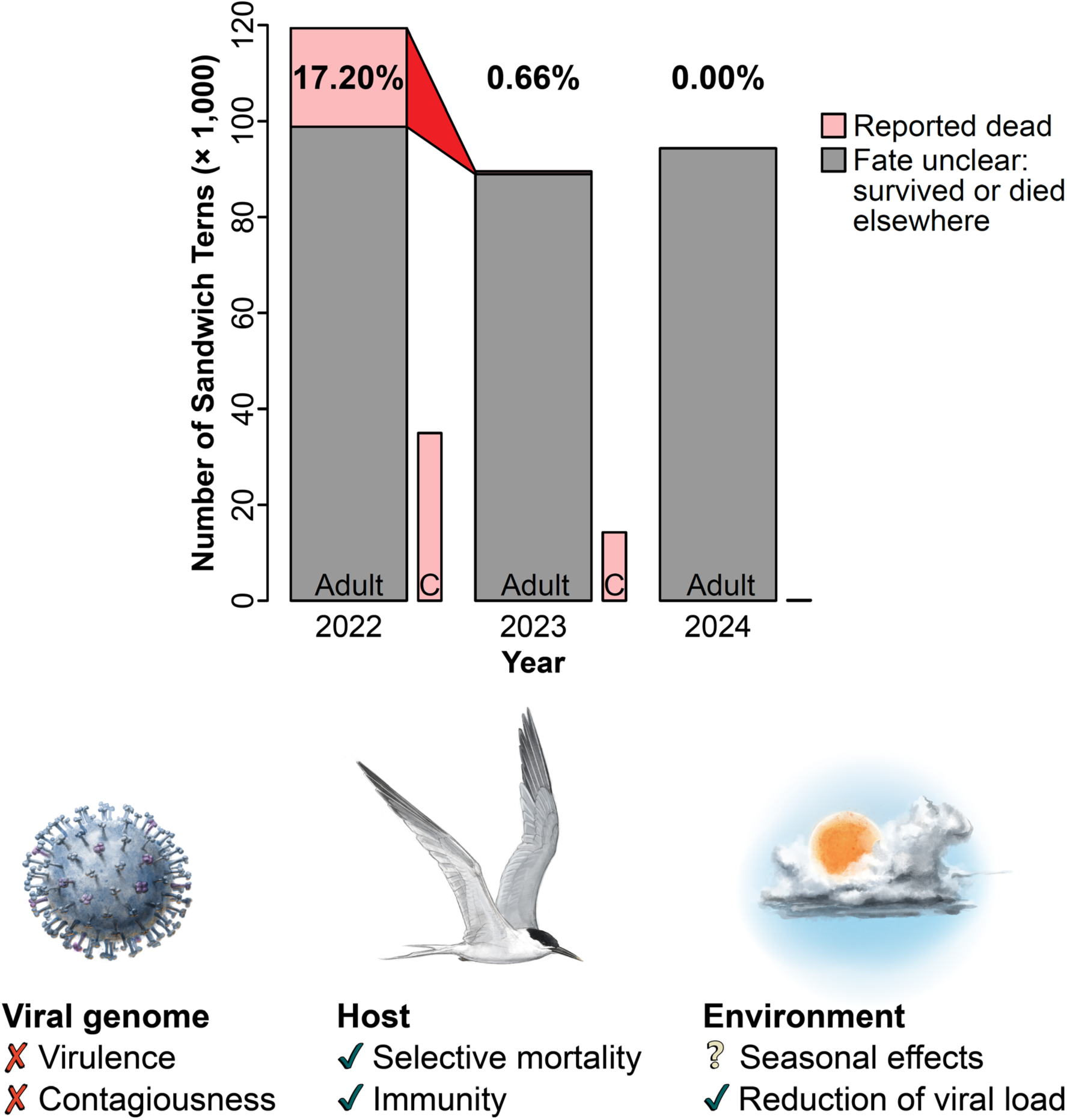
Breeding bird population size and mortality in the Northwestern European Sandwich Tern population during the three years since the HPAI H5N1 outbreak in 2022. Adult mortality is based on counts of dead birds found in colonies and along shorelines; chick mortality (abbreviated as “C”) is mostly estimated based on available reports of breeding success or estimates of chick deaths. The decline in mortality from 2022 to 2023 is highlighted, and possible explanations for this change are illustrated below. Either the virus, the host, or the environment must have changed. Available evidence suggests that the virus did not alter its virulence or contagiousness, but rather that host populations adapted through selective mortality and the build-up of immunity. Evidence for environmental effects is mixed: seasonal factors, such as rising temperatures or increased UV exposure, may have played a role, and environmental viral loads were likely reduced due to large-scale carcass removal adopted in 2023, and possibly also due to lower breeding bird densities in some species (e.g., Common Terns). The illustrations were created by Javier Lazaro.

Several factors may be at play. First, populations may have partially acquired population-level immunity through seroconversion of previously immunologically naïve adults surviving the initial outbreak. Around 20% of adult breeding Sandwich Terns in Germany tested positive for antibodies against H5Nx in both 2023 and 2024 (*N* = 50 in 2023 and *N* = 60 in 2024). However, it is important to note that birds with antibodies may still be susceptible to dying from the disease (Bouwhuis *et al*. in preparation), and it remains unclear whether these antibodies are neutralizing and whether they were present in and prior to 2022. Nonetheless, these findings raise the possibility that an increasing level of immunity helped protect affected populations from mass mortality. Consistently, Common Terns that were severely affected in both 2022 and 2023 in northern Germany showed a marked increase in H5Nx antibody prevalence, from less than 20% to around 40% between 2023 and 2024 (Knief *et al*. in preparation).

Additionally, the most susceptible individuals may have been removed from the population during the initial outbreak. Analyses of 29 years of European bird ringing and recovery data (1995–2023: 75,000 unique recoveries from 145,000 ringed birds) revealed that age classes were not equally affected by HPAI. Specifically, older birds appeared to suffer the highest mortality (Courtens *et al*. in preparation). This suggests that, by 2023, after the majority of the population had been exposed to the virus (Rijks *et al*. 2022), the most vulnerable individuals may have already been removed from the population or substantially reduced in number due to selective mortality (i.e., natural selection). Standing genetic variation at (host-specific) disease resilience genes may harbour adaptive variants that can rapidly rise in allele frequency under such extreme selection pressure. As a result, these individuals could no longer act as amplifier hosts and—naturally—also not contribute to the number of dead birds. Together, both immunity and selective mortality may have enhanced the protection of the remaining Northwestern European Sandwich Tern population.

Irrespective of the mechanism, the reduced adult mortality provided cautious optimism for population recovery, further reinforced by the absence of reported mass mortality events in Africa during the winter of 2023–2024. The specific virus variant (genotype EA-2022-BB), which was particularly well-adapted to gulls and caused severe mortality in Black-headed Gulls and Common Terns during the 2023 breeding season, disappeared in September 2023 (Fusaro *et al*. 2024). It was replaced by variants that seem to be less adapted to gulls and terns. In line with this, the breeding season of 2024 saw no noticeable infections or mortality among adult Sandwich Terns or their chicks, nor among other colonial breeding birds that had been heavily affected in previous years, such as Black-headed Gulls and Common Terns. The Northwestern European Sandwich Tern population remained at a reduced level, but increased substantially to 52,634 breeding pairs (Courtens *et al*. in preparation) (**Fig. 3**). This sudden rise may reflect the recruitment of previously non-breeding or younger individuals filling vacant breeding sites. These individuals may represent a “silent reserve” within the population. However, this reserve may now be largely depleted, suggesting that further recovery toward pre-HPAI population levels is likely to be slow and may take decades.

## 4. General Patterns in HPAI Outbreak Dynamics Among Wild Bird Species

### Anticipated Spread, Unpredictable Impact

The global spread of HPAI has been anticipated and discussed among researchers since its expansion out of Asia in 2005 (Butler & Ruttimann 2006), especially with regard to its potential arrival in South America and Antarctica (Global Consortium for H5N8 and Related Influenza Viruses 2016; Hurt *et al*. 2016). Rather than being unexpected, the scale of the current panzootic aligns with these earlier warnings. However, outbreaks have appeared idiosyncratic, meaning that it is next to impossible to predict which species will be affected next, where and when outbreaks will occur, or how severe they will be. Consequently, continuous monitoring and data collection remain essential (Caliendo *et al*. 2024; Nguyen *et al*. 2025).

### Varying Impact Across Species and Populations

The impact of HPAI varies considerably between species. For example, it has been devastating for the Great Skua (*Stercorarius skua*), a scavenging seabird species, with more than half of the global population lost (Tremlett *et al*. 2024). Some South American and South African seabird species suffered substantial losses as well, accounting for up to 50% of their world populations (Abolnik *et al*. 2023; Kuiken *et al*. 2025). Arguably, the most accurate data come from Northwestern European Sandwich Terns, where mortality exceeded 17% of the breeding bird population in 2022 (Knief *et al*. 2024), and a 25% loss in breeding pairs in 2023. Northern Gannets were affected throughout their global distribution range (Avery-Gomm *et al*. 2024; Lane *et al*. 2024), but reliable data on population-level impacts are lacking, though estimates suggest a loss of more than 10% of the world population. In contrast, Great Cormorants (*Phalacrocorax carbo*) in the Baltic Sea region were subject to limited outbreaks, which had little to no effect on population size (Bregnballe *et al*. 2024). Some species may even appear asymptomatic when infected with HPAI and thus serve as reservoirs. This includes Mallards (*Anas platyrhynchos*), some other dabbling ducks (Gaidet *et al*. 2008; Koethe *et al*. 2020; Teitelbaum *et al*. 2023), and possibly also larger gull species (*Larus* spp.) (Arnal *et al*. 2015; Tarasiuk *et al*. 2022).

Even closely related species may differ in their susceptibility to the disease. While mortality among secondary-host adult Bald Eagles (*Haliaeetus leucocephalus*) in North America was high, adults of the closely related European White-tailed Eagle (*Haliaeetus albicilla*) were not affected in Germany, despite likely being exposed to the virus, as indicated by the HPAI-related death of their chicks during the beginning of the enzootic in Europe (Günther *et al*. 2024; Nemeth *et al*. 2023). In Norway, however, some adult White-tailed Eagles were affected (Bøe *et al*. 2024), suggesting potential regional differences in susceptibility or exposure. Beyond these inter- and intraspecific differences, mortality rates also varied considerably among populations of other species. For example, during the 2022 breeding season in Germany, mortality ranged from 0% to nearly 50% in different Common Tern colonies and from 1% to over 60% in Sandwich Tern colonies (Pohlmann et al. 2023).

### Key Factors Influencing HPAI Dynamics

From this variation across species and populations, we can distil some factors influencing the dynamics of HPAI outbreaks that seem to be general across species:

i. Breeding bird density appears to be an important factor influencing both the risk and severity of outbreaks, as observed in Northern Gannets (Lane *et al*. 2024), Common Terns (Marchowski *et al*. 2024), and Great Cormorants (Bregnballe *et al*. 2024). Species-specific behaviours may further amplify transmission risk. For example, defecation within breeding colonies by Sandwich Terns, kleptoparasitism of gulls, frigatebirds and skuas (Gorta *et al*. 2024), or the use of communal club and bathing sites by Great Skuas, referred to as “ponds of death” due to their association with repeated mass mortality events (Bregnballe *et al*. 2023; Camphuysen & Gear 2022), may facilitate large-scale outbreaks. Common Terns have also been observed attacking sick, abnormally behaving conspecifics, which may also contribute to viral spread.
ii. Carcass removal can be an effective mitigation measure to reduce further spread of the virus (Knief *et al*. 2024). However, its success likely depends on early outbreak detection. Otherwise, it could potentially even have the opposite effect at the population level, if infected birds leave the colony in response to human disturbance and thereby spread the virus to other breeding sites.
iii. Instead of the virus altering its virulence or contagiousness, or the environment undergoing substantial changes, it appears that host populations may adjust to the virus over time through individual-level responses (phenotypic plasticity, adaptive immunity) and population-level processes (senescence-driven mortality, natural selection). The build-up of immunity and selective mortality of the most vulnerable individuals, i.e. those that previously acted as amplifier hosts, may have reduced exposure and provided greater protection for the remaining individuals (**Fig. 3**).
iv. Age-related mortality may arise from multiple, potentially interacting factors. In Sandwich Terns, immunosenescence (Bartleson *et al*. 2021) and age-dependent variation in breeding propensity may both contribute (Courtens *et al*. in preparation). Similar patterns have been observed in Common Terns, where differences in phenology or breeding status are linked to differential mortality risk (Bouwhuis *et al*. in preparation). In addition, genetic predisposition may contribute to selective mortality, favouring individuals with stronger immunity, or greater resistance or tolerance to infection (Chen *et al*. 2021). Further genomic studies in host species are needed to better understand the genetic basis of host-specific disease resilience and selective survival in wild birds.
v. Improved protection may be reflected by the fact that subsequent mass mortality events appear to become less likely after an initial severe outbreak. For instance, the outbreaks with the most profound impact in North America occurred in eastern Canada in 2022, resulting in the death of over 40,000 wild birds (Avery-Gomm *et al*. 2024). By 2023, the mortality rate in the region had dropped by 93%, and the few HPAI outbreaks that did occur primarily affected different species than in the previous year (Cormier *et al*. 2024). Similarly, observed mortality rates were 42% lower in 2023 compared to 2022 in Common Terns (Bouwhuis 2025) and no major outbreaks or mass mortalities were reported in adult Northern Gannets (Cormier *et al*. 2024; Tyndall *et al*. 2024), Sandwich Terns (Courtens *et al*. in preparation), or South American seabirds in 2023 either. The fact that large numbers of Sandwich Tern chicks died in 2023 may further suggest that adults were protected by acquired immunity, a hypothesis that should be validated through longitudinal seroprevalence studies.
vi. Seabirds typically exhibit delayed maturation, spending their early years away from the breeding grounds (Hamer *et al*. 2001). For example, Sandwich Terns remain in Africa for about three years before returning to Europe to breed for the first time (Veen 1977). These young birds could be considered a “silent reserve”, as they appear to fill vacant breeding sites—a pattern observed in Common Guillemots (*Uria aalge*) and Common Terns in 2023 and 2024 (Birkhead & Hatchwell 2025, Bouwhuis *et al*. in preparation), and which also seems to apply to Sandwich Terns.

### A Glimpse of Hope?

While the immediate impacts of HPAI have often been catastrophic for several seabird species, these last points suggest that some species may still have potential to recover, provided that populations retain sufficient numbers of individuals and that conservation efforts focus on long-term population resilience, protecting them from other potential (anthropogenic) threats.

## 5. Relevance of Quantitative Mortality Data Collection for Nature and Species Conservation

In today’s world, wildlife faces numerous threats, such as habitat loss, climate change and pollution, leading to a largely human-induced sixth mass extinction (Ceballos *et al*. 2015). Seabirds are the world’s most threatened group of birds (Paleczny *et al*. 2015) with many species facing various additional threats, including fishing, tourism, and the introduction of predators to previously suitable breeding sites, all contributing to their population declines worldwide (Phillips *et al*. 2023). The recent HPAI outbreaks must now be added to this growing list of threats. If HPAI continues to recur in seabird populations, it could become devastating for some species, fast.

### Implications for Bird Monitoring Programs

Due to their sensitivity, widespread distribution, and ease of observation, birds serve as prime bioindicators, offering insights into environmental conditions across different trophic levels. Consequently, numerous monitoring programs track avian population trends and investigate the causes of population changes. Without comprehensive mortality data from HPAI outbreaks, sudden population declines would remain unexplained. Such data are therefore vital for seabird monitoring and must be maintained, even during acute outbreaks in wild bird populations.

When combined with further monitoring efforts, long-term population studies facilitate more in-depth epidemiological analyses. For instance, the analysis of selective age-related mortality in Sandwich Terns was only possible due to extensive and continuous ringing efforts across Europe. Over the past 29 years, more than 145,000 Sandwich Terns were ringed, yielding almost 75,000 unique recoveries, which provided a robust foundation for survival modelling. Such efforts were initially aimed at understanding the natural history of the species, but have proven invaluable for addressing emerging conservation challenges (Johnston *et al*. 2025). They should, therefore, receive greater support and funding for other species as well (also see Clutton-Brock & Sheldon 2010).

### Implications for Conservation

In the case of Sandwich Terns, quantitative mortality data played a crucial role in prompting the German Lower Saxony Wadden Sea National Park administration and the German Federal Agency for Nature Conservation (BfN), together with the Federal Ministry for the Environment, Nature Conservation, Nuclear Safety and Consumer Protection (BMUV), to initiate a species conservation program. A key focus of this program is the establishment of effective preventive measures. These include an early warning system, currently implemented through video surveillance, and, in one colony, active surveillance. As soon as an abnormally behaving or dead bird is detected, testing can be initiated, and measures can be taken to contain the infection. Active surveillance, i.e. monitoring the prevalence of HPAI viruses in Sandwich Terns, is carried out by capturing breeding birds and testing them for the presence of viral RNA using molecular methods. Such measures should also be implemented for other species that have suffered similarly from recent HPAI outbreaks.

As breeding bird density is a key factor influencing both the risk and severity of outbreaks, reducing densities or the number of birds in a colony could lower infection risk and limit outbreak sizes. One potential strategy is the creation or restoration of natural breeding habitats. While this may be challenging for species such as Sandwich Terns (though still possible, for example through the expansion of suitable protected areas), it may be more readily achievable for other species. For instance, Common Terns readily nest on floating platforms (e.g. Manikowska–Ślepowrońska *et al*. 2022), and increasing the number of these platforms could help distribute birds more evenly, thereby potentially reducing individual densities and the risk of disease transmission.

### Implications for Vaccination

Particularly vulnerable species, such as the African Penguin (*Spheniscus demersus*) in South Africa (Roberts *et al*. 2024) and the California Condor (*Gymnogyps californianus*) in the USA (U.S. Fish & Wildlife Service 2024), have been vaccinated against HPAI. A trial involving 24 captive African Penguins tested different vaccine types and strategies (with or without booster) and induced antibody titres expected to protect from virus shedding and mortality. So far, the strength of the immune response and the duration of protection varied depending on the vaccine type and strategy, but protection never lasted beyond a single breeding season, such that further study will be necessary.

After 21 of the remaining 561 California Condors died from HPAI in early 2023, the surviving individuals were vaccinated under strict regulations. By October 2024, nearly half of the population (*N* = 207) had received at least one dose, and no further casualties were reported among them.

For a species like the Sandwich Tern, which is fortunately still relatively common, such measures are practically impossible and politically undesired. Moreover, as an RNA virus with a segmented genome, HPAI viruses are highly mutable, facilitating the emergence of new variants that may render existing vaccines ineffective, necessitating continuous re-vaccination and comprehensive surveillance of wild bird populations to detect and monitor potential escape mutants.

### Shifting Focus from Prevention to Early Detection and Response

Since HPAI cannot be eradicated in wild populations, management efforts must, at least at present, focus on early detection and rapid response rather than prevention. Vaccination may offer a solution for selected, highly endangered species, but it likely remains impractical for most wild birds, particularly in the absence of non-invasive, passively delivered vaccination strategies that are capable of inducing sterilizing immunity. Likewise, controlling virus entry points seems infeasible given the complexity of transmission pathways. Given these constraints, the most effective strategy is to improve how we monitor diseases and respond to outbreaks, which hinges on robust, quantitative mortality data. Without such data, long-term conservation measures will remain dangerously inadequate.

## 6. Improving Quantitative Mortality Data Collection in Wild Animals

### The Need for Systematic Wildlife Mortality Monitoring

Current surveillance efforts in wild birds primarily focus on detecting the presence of the virus (EMPRES-i+), often through opportunistic sampling of sick or dead individuals, which is often limited to places where people occur in high densities. Although informative, such qualitative data yield only limited understanding of the true extent of mortality and its long-term consequences for affected populations (**Fig. 2**). To move beyond a reactive crisis response, we need a supportive, systematic and standardized approach to collecting quantitative mortality data, combined with diagnostics and genomic epidemiology (**Table 1**; Atkinson & Baillie 2025; Klaassen & Wille 2023). Coordinated bird and disease monitoring programs, improved data-sharing frameworks, and closer integration of ecological and epidemiological research are essential. These efforts would not only help quantify impacts more accurately, but also enable the development of an effective and robust early warning system for future outbreaks (Couty *et al*. 2025). The Pathogen Harmonized Observatory (PHAROS) database (pharos.viralemergence.org) is a promising global initiative but is still in its infancy (Schwantes *et al*. 2024).

**Table 1.**
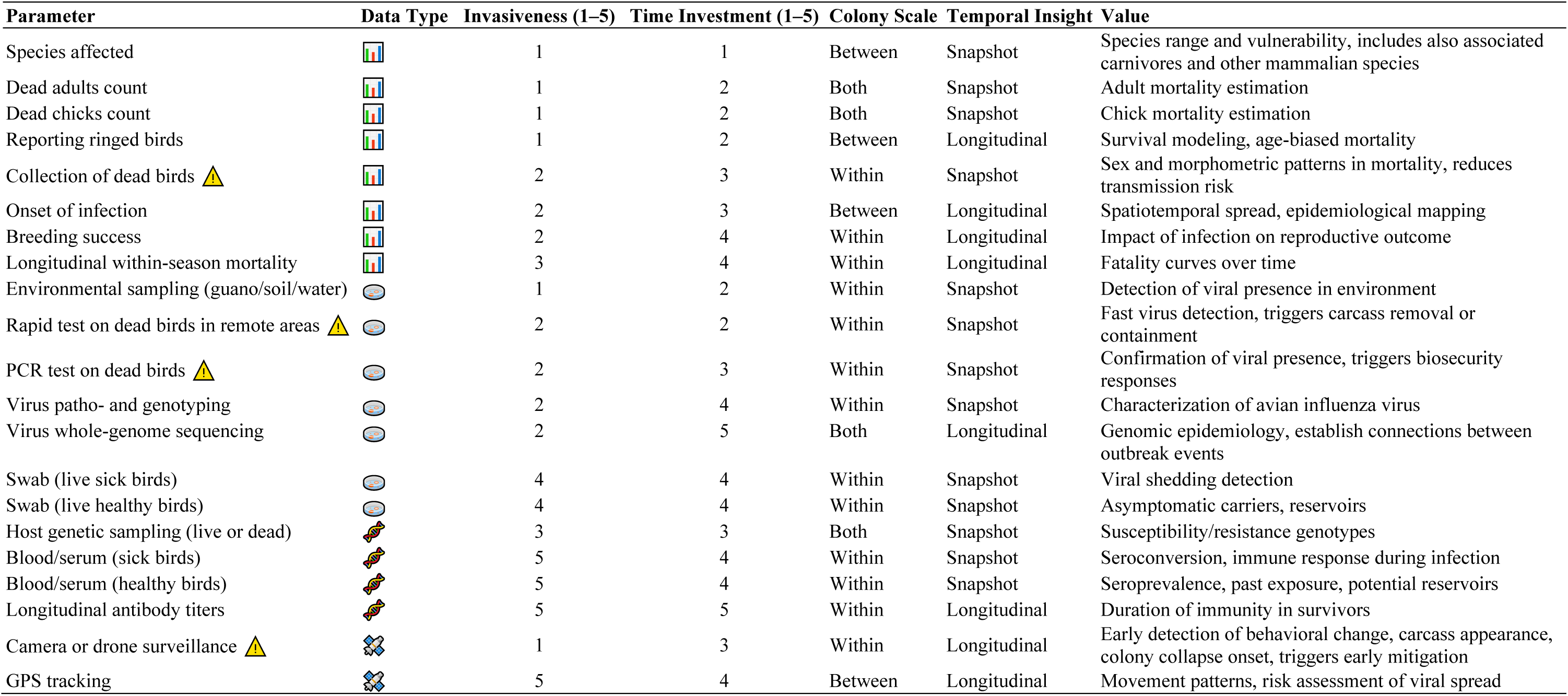
Overview of monitoring parameters for assessing the impact and dynamics of HPAI in seabird colonies. Parameters are classified by data type, invasiveness, time investment, major spatial inference scale (colony scale), and whether they provide snapshot or longitudinal insight. Invasiveness and time investment were assessed qualitatively based on field experience and expert judgement. The final column summarizes the primary value or application of each parameter in the context of HPAI surveillance. *Parameters* flagged with 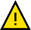 directly support mitigation measures during HPAI outbreaks. *Data type*: 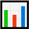 = Demographic or trait-based monitoring, 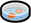 = Diagnostics / virological testing, 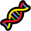 = Genetic or immunological insights, 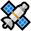 = Remote sensing / automated monitoring. *Colony Scale*: Within = intra-colony level understanding, Between = inter-colony level understanding, Both = intra- and inter-colony level understanding. *Temporal Insight*: Snapshot = One-time or point-in-time data, Longitudinal = Repeated measures needed.

For poultry, retrospective mortality data are accessible through systems such as animal disease insurance funds. In contrast, wildlife mortality data collection has been largely *ad hoc*, depending on private initiatives, existing records and contributions of local conservation managers, who themselves are not evenly distributed across wildlife populations. Until now, there has been no centralized system for systematically tracking wildlife mortality. However, the unprecedented scale of HPAI cases in wild birds since 2022 has made efforts to create such a system increasingly urgent.

A dedicated institution or program for systematic wildlife mortality monitoring would need to coordinate with relevant authorities, train personnel in biosafety and biosecurity, and develop standardized protocols for field data collection. Additionally, such an institution could expand and coordinate active surveillance, an approach increasingly recognized as essential by the European Union and global health organizations such as the World Health Organization (WHO), the Food and Agriculture Organization of the UN (FAO), and the World Organization for Animal Health (WOAH).

The World Animal Health Information System (WAHIS) of the WOAH collects data on epidemiologically important diseases in domestic and wild animals, including HPAI. It is the most comprehensive—and currently the only—global source of HPAI-related case numbers in wild animals (Klaassen & Wille 2023). However, the database is incomplete: numerous outbreaks are missing, and existing entries are often inconsistent and lack essential details. Reports frequently group multiple species within a single outbreak and do not specify whether adults or chicks were affected, making it impossible to reliably quantify species-specific impacts. Compared to our literature review of reported outbreaks in wild birds since September 2020, WAHIS data reflect a broadly similar geographic pattern, with case numbers per country correlating. Nevertheless, WAHIS underestimates total case numbers by one order of magnitude (**Fig. 2**).

Some regional systems do exist though: in the USA, the Wildlife Health Information Sharing Partnership (WHISPers) collects quantitative information on wildlife mortality events, but it suffers from the same lack of detail as the WAHIS database. In the United Kingdom, a smartphone app (Epicollect5) has been developed to streamline data entry and reporting. In response to the arrival of HPAI in the (Sub-)Antarctic region, the Scientific Committee on Antarctic Research (SCAR) has proposed innovative surveillance strategies and established an HPAI database to compile all suspected and confirmed outbreaks in the area, with the aim of monitoring the spread of the virus and better understanding its impacts on wildlife (Wille *et al*. 2025).

### Reduce Regulatory Barriers to Active Surveillance

Passive surveillance remains the most effective method for early detection of HPAI cases. However, it should be complemented by serological surveillance to monitor antibody levels over time and to better understand population-level exposure dynamics. Active surveillance can provide valuable data, particularly for research purposes. However, from an animal health perspective, maintaining such systems is often challenging due to the high logistical effort involved and the typically low virus detection rates. These constraints, combined with the fact that serological data are even scarcer than virus detection data, highlight the need for more integrated and sustained surveillance strategies that include serology as a core component in active sampling.

In some European countries, rigid and time-consuming animal welfare approval processes hinder both responsive and proactive surveillance. These bureaucratic hurdles can make timely investigations almost impossible, especially given the unpredictable nature of HPAI outbreaks. We often do not know in advance which species will be affected next. In contrast, some countries have more flexible regulatory frameworks, enabling scientists and authorities to respond faster and more effectively.

### Balance Data Collection with Biosafety and Biosecurity

Data collection during active outbreaks is essential for understanding transmission dynamics and for assessing population-level impacts. Delayed access to affected sites, whether due to regulations, logistics, institutional hesitancy, or concerns over welfare, can result in the loss of critical information. While the need to minimize disturbance and ensure biosafety is paramount, these goals can be met through the use of personal protective equipment (PPE), strict field protocols, and collaboration with veterinary authorities (Bregnballe *et al*. 2023; Nature Medicine Editorial 2024). To improve outbreak responses, site managers and government administrations must find a workable balance between data collection needs and animal and human welfare.

### Bridging the Gap Between Field and Laboratory Data

Quantitative field data and qualitative laboratory data must be better integrated. The two data sources are generated using different methodologies, techniques, and risk assessments, which can create challenges in combining them effectively. Differences in sampling effort, detection probabilities, and diagnostic sensitivities further complicate direct comparisons. Inconsistent communication and limited mutual understanding between field and laboratory teams can also result in mismatched expectations and data gaps. Addressing these challenges requires standardized protocols and interdisciplinary collaboration. Building stronger networks and fostering mutual trust between ecologists, epidemiologists, and laboratory researchers are essential to ensure responsible data collection and accurate risk assessments. This could be realized by regular subject-specific cross-sectoral meetings and feedback loops, interdisciplinary workshops, shared digital platforms and databases, and long-term institutional partnerships formalized through memoranda of understanding (MoUs) between academic institutions, public health agencies and governmental institutions.

### Increase Funding and Institutional Support

As the global spread of HPAI is likely to continue, delaying the establishment of structured monitoring systems will only weaken the ability to respond to future outbreaks. Yet, securing funding for interdisciplinary research remains a major hurdle.

Despite the urgency of recent outbreaks, researchers and institutions attempting to implement integrated surveillance—combining ecological analysis, genetic research, and field-based data collection—have often struggled to secure adequate support. Proposals are frequently deemed insufficiently “medical”, not focused enough on direct species conservation, or not interdisciplinary in the narrow ways defined by funding agencies. These recurring obstacles point to a deeper issue: the importance of integrated wildlife mortality data collection is still not fully recognized within existing funding frameworks.

This mindset must change. Rigid frameworks and siloed thinking are no longer fit for purpose in the face of complex health challenges. New concepts and approaches have already been proposed (e.g. Kuiken 2021), but their implementation remains slow. Without more flexible funding mechanisms and stronger institutional support, our capacity to respond to future emerging disease outbreaks will remain reactive rather than proactive, which places both wildlife and public health at continued risk. The European Wildlife Disease Association (EWDA) is trying to address this by promoting transformative research in wildlife health. However, meaningful progress requires a stronger and sustained commitment from national governments and funding agencies. Only with coordinated investment in long-term infrastructure, workforce development, and cross-sectoral research can we build a truly resilient One Health approach.

## 7. Quantitative Wildlife Mortality Data is Essential for the One Health Approach to HPAI

Effectively containing HPAI requires a comprehensive understanding of the interplay between the virus, the environment, and its hosts. A One Health approach (Bertram *et al*. 2024; Harvey *et al*. 2023) is essential, one that explicitly integrates wildlife into surveillance and response efforts, even as poultry farming remains the primary economic focus.

### HPAI Transmission Is a Two-Way Street

The transmission of HPAI between wild birds and poultry is not unidirectional. While poultry farms often act as the source of infection (Kuiken & Cromie 2022), wild birds can also introduce the virus to commercial flocks and even mammalian populations. For example, HPAI likely reached North America via wild birds from Iceland and Greenland (Ahrens *et al*. 2024; Caliendo *et al*. 2022; Günther *et al*. 2022; Harvey *et al*. 2023) and later arrived in Hawai‘i via migratory birds traveling from Alaska along the Pacific flyway (State of Hawai’i 2025). Moreover, reports from the European Food Safety Authority (EFSA) document numerous instances of wild birds transmitting the virus to both poultry and mammalian farms across Europe.

When large numbers of wild birds become infected, the viral load in the environment rises, increasing the risk of transmission not only to poultry, but also to mammals, including humans. Understanding and managing HPAI in wildlife is therefore not just a matter of species conservation, it has far-reaching economic and public health implications.

### Ethical Responsibility

Humans have played a direct role not only in causing the decline of many species, but also in the emergence of HPAI, as these viruses evolved in intensive poultry farming. Wild birds are not the cause of this crisis. They are its victims. Some species may never fully recover from these outbreaks. A critical step in addressing this crisis is the systematic collection of quantitative wildlife mortality data. Such data are vital for moving from reactive, short-term crisis management to proactive evidence-based prevention, conservation and long-term containment strategies. Ultimately, it is our ethical responsibility to act, not only to conserve wildlife and maintain ecosystem stability, but also to safeguard human health and prevent future zoonotic threats.

## Conclusions

- The emergence and global spread of HPAI H5N1 clade 2.3.4.4b in wild bird populations has exposed critical gaps in our understanding of the dynamics and impacts of HPAI outbreaks in wild animal populations (Couty *et al*. 2025; Klaassen & Wille 2023). While the devastating consequences are qualitatively evident, we remain unable to provide accurate and precise assessments of the ecological impacts for most species and ecosystems. This shortcoming arises because the currently established global HPAI reporting systems were not designed with wildlife in mind. Although they provide valuable information, these general-purpose disease surveillance tools are primarily intended for domestic livestock. As a result, they are poorly suited to capture the scale, species-specific impacts, and ecological consequences of HPAI outbreaks in wild populations.
- We urgently need dedicated and systematic wildlife mortality monitoring, which includes innovative surveillance strategies for remote regions, streamlined data entry solutions (Wille *et al*. 2025), and open-access data sharing (Schwantes *et al*. 2024). Such a system must integrate the mortality data with diagnostic and viral genetic data, which typically become available at different times during an outbreak. The database established by the Scientific Committee on Antarctic Research for monitoring HPAI in Antarctic wildlife may serve as a promising model. Combined with, for example, smartphone-based reporting tools, such a system could set a new gold standard for global wildlife mortality data collection.
- Addressing this data deficiency is not merely a scientific necessity, but an ethical imperative. Wild birds are an integral part of the current HPAI pandemic, but they are not its origin. They are suffering the consequences of a crisis largely driven by human agricultural practices. Even with limited data and resources, research during the current pandemic has yielded important insights into viral evolution, host adaptation, and outbreak mitigation in wildlife. Critically, it has also revealed that host species characteristics—such as immunity, behaviour, and ecology—play a fundamental role in shaping outbreak dynamics and outcomes.
- What we now need is recognition that HPAI in wildlife is not only a conservation issue threatening biodiversity, but also a serious concern for public health and food security. We call for political commitment to support ecological research as an integral part of safeguarding against the current and future zoonotic threats.

## Supporting information

References for Table S1

## Acknowledgements

We are grateful to Dana Thal (One Health Platform) for inspiring the thought process that led to this perspective. We thank Javier Lazaro for his illustrations and Martin Beer for thoughtful and valuable feedback on an earlier version of this manuscript.

## Author Contributions

Conceptualization and visualizations: UK. Writing: UK prepared the original draft. The final manuscript incorporated input from all authors, and all authors approved the final version before submission.

## Competing interests

The authors declare no competing interests.

## Additional Information

*Supplementary Table 1*: Quantitative HPAI H5N1-related mortality data in wild animals, extracted from the literature since September 2020.

## Notes

### Competing Interest Statement

The authors have declared no competing interest.

