## Supplementary material for "Counting Cases, Conserving Species: Addressing Highly Pathogenic Avian Influenza in Wildlife": References for Table S1

**Ulrich Knief[1,*], Sandra Bouwhuis[2], Anja Globig[3], Anne Günther[4], Wouter Courtens[5]**

[1] University of Freiburg, Institute of Biology I (Zoology), Evolutionary Biology and Ecology, Hauptstr. 1, DE-79104 Freiburg, Germany. Orcid: 0000-0001-6959-3033

[2] Institute of Avian Research, An der Vogelwarte 21, D-26386 Wilhelmshaven, Germany. Orcid: 0000-0003-4023-1578

[3] Friedrich-Loeffler-Institut, Institute of International Animal Health/One Health, Federal Research Institute for Animal Health, Südufer 10, DE-17493 Greifswald-Insel Riems, Germany. Orcid: 0009-0006-1966-6760

[4] Friedrich-Loeffler-Institut, Institute of Diagnostic Virology, Federal Research Institute for Animal Health, Südufer 10, DE-17493 Greifswald-Insel Riems, Germany. Orcid: 0000-0001-5176-6346

[5] Research Institute for Nature and Forest, Havenlaan 88, bus 73, 1000 Brussels, Belgium. Orcid: 0000-0002-5806-0343

[*] Address for correspondence: Ulrich Knief, University of Freiburg, Institute of Biology I (Zoology), Evolutionary Biology and Ecology, Hauptstr. 1, DE-79104 Freiburg, Germany, Phone: 0049-761-203-2911,
